# PFOS Disrupts Membrane Signaling and Epithelial Integrity in Fallopian Tube Cells

**DOI:** 10.64898/2026.03.24.713959

**Authors:** Tonja Pavlovič, Sadaf Farsinejad, Dibyendu Sarkar, Benjamin Tycko, Marcin Iwanicki

## Abstract

Perfluorooctane sulfonic acid (PFOS), a per- and polyfluoroalkyl substance (PFAS), is a widespread persistent environmental pollutant that has been implicated in various human health conditions, including infertility and cancer. Here, we investigate the effects of acute exposure to PFOS on human fallopian tube epithelial (FNE) cells that are essential for fertility and increasingly recognized as the origin site for high-grade serous ovarian cancer. We show that acute PFOS exposure changes morphology, arrests proliferation, impairs adhesion, and compromises epithelial integrity of FNE cells. Using transcriptomic profiling of FNE cells exposed to PFOS, we found increased expression of genes associated with stress-response signal transduction, including KRAS, and decreased expression of genes related to cholesterol transport and lipid homeostasis. We show that inhibition of MEK/ERK or cholesterol supplementation rescued changes in cell morphology. Further, we performed membrane fluidity measurements of cells exposed to PFOS and found elevated membrane disorder and fluidity. Our results are consistent with a model in which PFOS perturbs plasma membrane, activates stress-response signaling pathways, and impairs epithelial cell function. These studies establish a framework for understanding the effects of PFAS on cell physiology.

## INTRODUCTION

Perfluorooctane sulfonic acid (PFOS) is a synthetic organofluorine compound that is part of a broader class of per- and polyfluoroalkyl substances (PFAS). Due to its exceptional water-repellent properties, PFOS has been extensively utilized in industrial and consumer products like firefighting foams, textiles, food packaging, and intimate care products (Buck et al., 2011).

PFOS is environmentally persistent and can bioaccumulate (Buck et al., 2011, Giesy and Kannan, 2002). It has been detected in a myriad of natural materials, from soil and groundwater to human tissues (Buck et al., 2011). Reported concentrations of PFOS in blood serum vary greatly, with the average being in the nanomolar range (20–80 nM) in nonoccupational adult populations, and higher levels in occupationally exposed individuals (up to 3 μM) (Olsen et al., 2003a, Olsen et al., 2003b, Olsen et al., 2003c). Importantly, PFOS is one of the most abundant PFAS that accumulates in reproductive organs and fluids, with concentrations in follicular fluid reported in the nanomolar range (2–60 nM) (Zeng et al., 2023, Petro et al., 2014, Kim et al., 2020). Despite low-level exposures, physiological concentrations can become significant due to bioaccumulation and food chain effects (Giesy and Kannan, 2002). In addition, indirect measurements of PFOS in bodily fluids may be unreliable because PFOS binds strongly both in vivo with serum proteins and in vitro with plastic tubes and glassware, underscoring the variability in concentrations reported across studies (Zhang et al., 2009, Lath et al., 2019).

Nevertheless, numerous epidemiological studies have clearly linked PFOS exposure to a range of health issues, including thyroid dysfunction, ulcerative colitis, testicular and kidney cancer, hypertension, and infertility (Steenland et al., 2010, Lopez-Espinosa et al., 2012, Joensen et al., 2009, Steenland et al., 2013, Fart et al., 2021). However, the impact of PFOS on the human fallopian tube epithelium remains poorly understood. This tissue plays a vital role in egg transport, fertilization, and early embryonic development (Li and Winuthayanon, 2017, Coy and Yanagimachi, 2015). Moreover, fallopian tube epithelium is increasingly recognized to be the tissue of origin for high-grade serous ovarian carcinoma (Perets and Drapkin, 2016). In particular, precursor lesions known as serous tubal intraepithelial carcinomas (STICs) arise within this epithelium and represent the earliest morphologically identifiable stage of this lethal cancer (Ducie et al., 2017, Li et al., 2014). This underscores the critical role of fallopian tube epithelial health in both reproductive biology and disease.

Beyond its broad effects on health, PFOS has been consistently linked to alterations in lipid metabolism and direct interactions with cellular membranes (Hu et al., 2003, Zhao et al., 2023). In animal studies, PFOS exposure reduced circulating lipids and lipoproteins while altering hepatic expression of genes involved in fatty acid synthesis, transport, and oxidation (Wang et al., 2014). In human hepatocyte-like models, PFOS induced triglyceride accumulation and dysregulated PPARα-target gene expression, suggesting direct interference with lipid and cholesterol homeostasis (Sadrabadi et al., 2024, Kashobwe et al., 2024). Consistent with these findings, epidemiological analyses have reported associations between PFOS levels and circulating cholesterol (Haug et al., 2023). At the cellular level, PFOS exhibits a strong affinity for lipid assemblies, readily partitioning into phospholipid bilayers and integrating into cell membranes, with potential consequences for membrane order and receptor function (Xie et al., 2010, Hu et al., 2003). These findings provide a mechanistic basis for how PFOS could perturb epithelial cell physiology, including the fallopian tube.

To investigate the effects of PFOS on fallopian tube epithelium, we used immortalized human fallopian tube non-ciliated epithelial cell model (FNE). We observed that PFOS alters FNE cell shape, adhesion, proliferation, motility, and epithelial integrity. Gene-expression profiling indicated PFOS-mediated induction of stress-response pathways, such as KRAS/MAPK signaling, as well as changes in cholesterol transport and lipid homeostasis. While agonists or antagonists of PPARα receptor, a nuclear transcriptional regulator that activates genes involved in lipid metabolism, failed to restore PFOS-induced cell shape defects, pharmacologic blockade of mitogen-activated protein kinase kinase (MEK) largely rescued cells from this PFOS-induced phenotype.

We hypothesized that activation of membrane-proximal signaling pathways, such as MAPK, may be linked to a disruption in membrane homeostasis. As PFOS is known to be able to insert into the plasma membrane, we tested the idea that PFOS insertion into the plasma membrane, at a critical concentration, could perturb the plasma membrane and lead to a downstream signaling response. This was supported by membrane fluidity measurements, which showed that PFOS increases membrane disorder and fluidity. Additionally, cholesterol supplementation restored FNE cell shape, membrane order and fluidity, and abrogated cytotoxicity associated with increasing PFOS exposure. Our findings suggest that exposure of FNE cells to PFOS disrupts membrane integrity and may lead to altered receptor signaling, resulting in activation of stress response, loss of cell adhesion, and impairment of epithelial integrity.

## RESULTS

### PFOS alters cell morphology

To test the effect of PFOS acute treatment, we treated FNE cells with a single dose of 25 μM PFOS for a period ranging from 48 to 96 hours. Phase contrast imaging revealed a striking change in cell morphology compared to vehicle-treated control (**Figure 1A**). FNE cells exposed to PFOS exhibited less circular and a more elongated, spindle-like shape (**Figure 1A, S1A**). This morphological transition was not unique to wild-type FNE cells. To varying degrees, shape alterations upon acute exposure to PFOS were detected in multiple cell lines tested (**Figure 1A**). Similar to wild-type FNE cells, dramatic shape changes were observed in FNE cells expressing mutant p53 (FNE ΔTP53) and immortalized retinal pigmented epithelium (RPE1) cells. Three distinct ovarian carcinoma cell lines with epithelial-like features, CaOV3, Kuramochi, and RMUGS, responded to acute PFOS treatment to a variable extent, with CaOV3 changing their shape less drastically than Kuramochi and RMUGS (**Figure 1A**). Quantitative image analysis confirmed an increase in length versus width ratio in majority of tested cell lines (**Figure 1B**) and a significant decrease in FNE cell circularity index (**Figure S1A**). The observed changes in cell shape appeared to be contact-dependent and unlikely to result directly from a decrease in cell size as we could only observe a marginal reduction in suspended cell volume across multiple cell types following PFOS exposure, which was not significant enough to fully account for the striking shape alterations when the cells were adhered (**Figure 1C**). Notably, the cell viability half-maximal inhibitory concentration (IC_50_) varied between cell lines, with FNE (IC_50_ = 57.47 ± 14.31 μM), FNE ΔTP53 (IC_50_ = 79.11 ± 61.55 μM), CaOV3 (IC_50_ = 46.12 ± μM), and RMUGS (IC_50_ = 15.87 ± μM) seemingly much more sensitive to PFOS, than RPE1 (IC_50_ = 1.37 ± 0.56 mM) and Kuramochi (IC_50_ = 2.47 ± mM) respectively (**Figure S1B**). In addition to internal variations in IC_50_ values between cell lines, we found that other external factors can substantially influence the efficacy of PFOS. One of the most prominent factors affecting PFOS efficacy was fetal bovine serum (FBS), a key supplement in cell culture media. Standard concentrations of FBS, widely used for *in vitro* cell culture support (5% and 10%), were able to fully abrogate the effect of PFOS on FNE cell morphology and resulted in a loss of cytotoxicity (IC_50_ = ∞; **Figure S1C**). This strongly suggests the presence of factors in FBS that can either directly affect PFOS bioavailability in solution (e.g., albumin) or indirectly impede its effects on FNE cells. Without FBS supplementation, FNE cells were significantly more vulnerable to PFOS exposure, with an IC_50_ value close to the concentrations previously reported for occupational exposure (IC_50_ = 11.61 μM; **Figure S1C**) (Olsen et al., 2003c). We conducted experiments by supplementing media with 2% FBS as a compromise between PFOS efficacy and supporting *in vitro* cell growth. Exposure of FNE cells to 25 µM PFOS resulted in increased PFOS levels detected in the cells, consistent with intracellular accumulation (**Figure S1D**).

**Figure 1.**
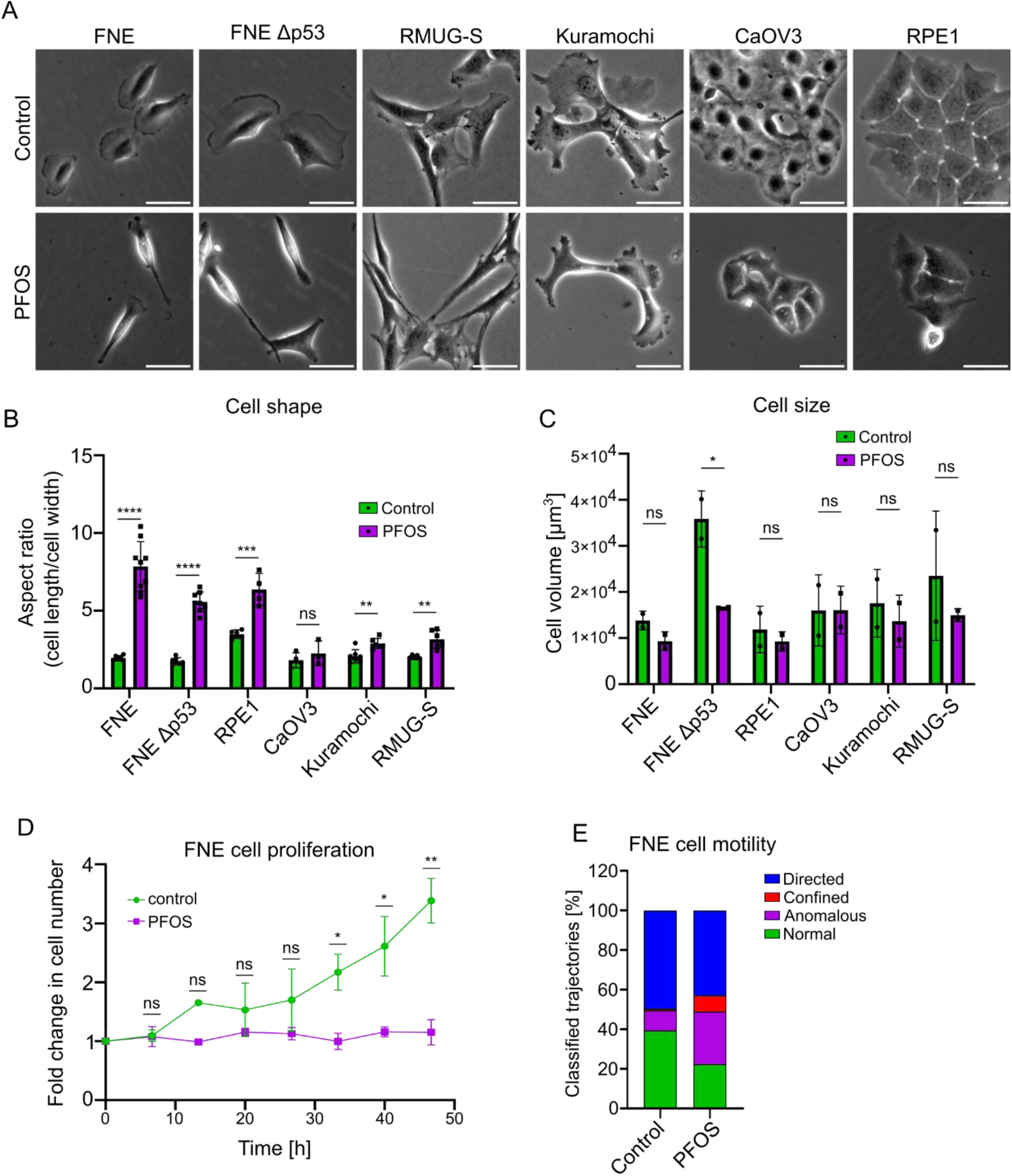
PFOS induces morphological changes in epithelial cells. **(A)** Phase-contrast images of normal fallopian tube epithelial cells (FNE), FNE cells expressing mutant p53 (FNE Δp53), immortalized retinal pigmented epithelial cells (RPE1), and ovarian carcinoma cell lines (CaOV3, Kuramochi, RMUG-S) 48 h after treatment with 25 µM PFOS or vehicle control (water). Images were acquired using a 10× objective. Scale bars = 50 µm. **(B)** Quantification of individual cell length-to-width (aspect) ratios for each cell line after 48 h exposure to PFOS or control. Each dot represents the mean of > 50 cells from one biological replicate (n ≥ 3). **(C)** Mean suspended-cell volume quantified by flow cytometry forward-scatter area (FSC-A) and converted to absolute cell volume using size calibration beads and linear regression. Each dot represents an independent repeat containing ≥ 1000 events each. **(D)** Live-cell imaging of FNE cell proliferation starting 48 h after PFOS or control treatment. GFP-positive cells were counted every 12 h across three fields of view per sample (n = 3). Data are shown as fold change relative to t = 0 h. **(E)** Representative classification of 24-h single-cell trajectories from time-lapse recordings into four motility categories: directed/active, confined, anomalous sub-diffusive, and normal diffusive motion. Bar segments show the average fraction (%) of each trajectory type compiled from ≥ 3 regions of interest per replicate (n = 3). All quantitative data are mean ± SD. Statistical comparisons were performed by unpaired t-test or two-way ANOVA with Sidak’s post hoc test unless otherwise indicated. *p < 0.05; **p < 0.01; ***p < 0.001; ****p < 0.0001; ns, not significant.

### PFOS arrests cell proliferation and impairs motility

Acute PFOS exposure led to a marked suppression of FNE cell proliferation, as demonstrated by live-cell imaging of confluency dynamics over time (**Figure 1D**). This inhibitory effect on proliferation was further validated by direct cell counting, which confirmed a significant reduction in FNE cell number following 192 hours of PFOS treatment compared to untreated FNE cells (**Figure S2A**). Interestingly, the effect of PFOS on proliferation and morphological changes was found to be largely reversible within 24 to 48 hours of PFOS washout indicating the possibility that PFOS-induced changes require sustained exposure (**Figure S2B,C**).

In addition to shape changes, PFOS-treated FNE cells exhibited distinct changes in migratory behavior. While PFOS treatment did not seem to slow them down, the overall distance that the PFOS-exposed FNE cells had travelled was shorter and closer to their initial starting point (**Figure S2D,E**; **Movie 1**,**2**). Detailed analysis of cell trajectories revealed a transition from mainly normal or actively directional motility in vehicle-treated control cells to a more confined or sub-diffusive motion in PFOS-exposed cells (**Figure 1E**). Subtle shifts in migratory dynamics can often indicate altered cytoskeletal coordination and possibly changes in how cells interact with their substrate.

To assess whether this effect was unique to PFOS or shared among related PFAS substances, we treated FNE cells with structurally similar PFAS analogs. Perfluoro<u>h</u>e<u>x</u>anoic <u>a</u>cid (PFHxA) induced only mild morphological changes under equivalent conditions, whereas the longer-chain compound <u>p</u>erfluoro<u>d</u>ecanoic <u>a</u>cid (PFDA) caused a pronounced elongation phenotype similar to, and in some cases more severe than PFOS exposure (**Figure S2F,G**). These results suggest that cell-shape alterations could correlate with PFAS chain length and hydrophobicity.

Taken together, the observed changes in contact-dependent cell morphology, proliferative capacity, and motility indicated that PFOS may disrupt cellular adhesion and mechanical interactions with the extracellular environment. These findings prompted a focused investigation into PFOS-induced changes in cell adhesion patterns and associated molecular pathways.

### PFOS reduces cell-substrate adhesion

Changes in cell shape and motility are often tied to cell adhesion patterns, as these properties are linked to the cytoskeleton and focal adhesion turnover (Parsons et al., 2010, Geiger et al., 2009). As expected, exposure to PFOS significantly decreased cell-substrate adhesion, marked by a reduction in the number of focal adhesions (**Figure 2A,B**). Whereas vehicle-control cells showed robust actin stress fibers, PFOS-treated cells largely lacked any discernible fiber-like structures (**Figure 2A, zoom box**). When FNE cells were detached briefly and then allowed to reattach, PFOS-treatment showed inefficient reattachment and delayed spreading compared to control cells (**Figure 2C**). Since PFOS-induced shape change appeared adhesion-dependent, we further tested whether enforcing stronger substrate attachment could restore normal morphology. Increasing substrate adhesion by fibronectin, laminin, or poly-L-lysine coating did not restore normal morphology (**Figure S3A**), suggesting that PFOS suppresses formation of cell substrate adhesion.

**Figure 2.**
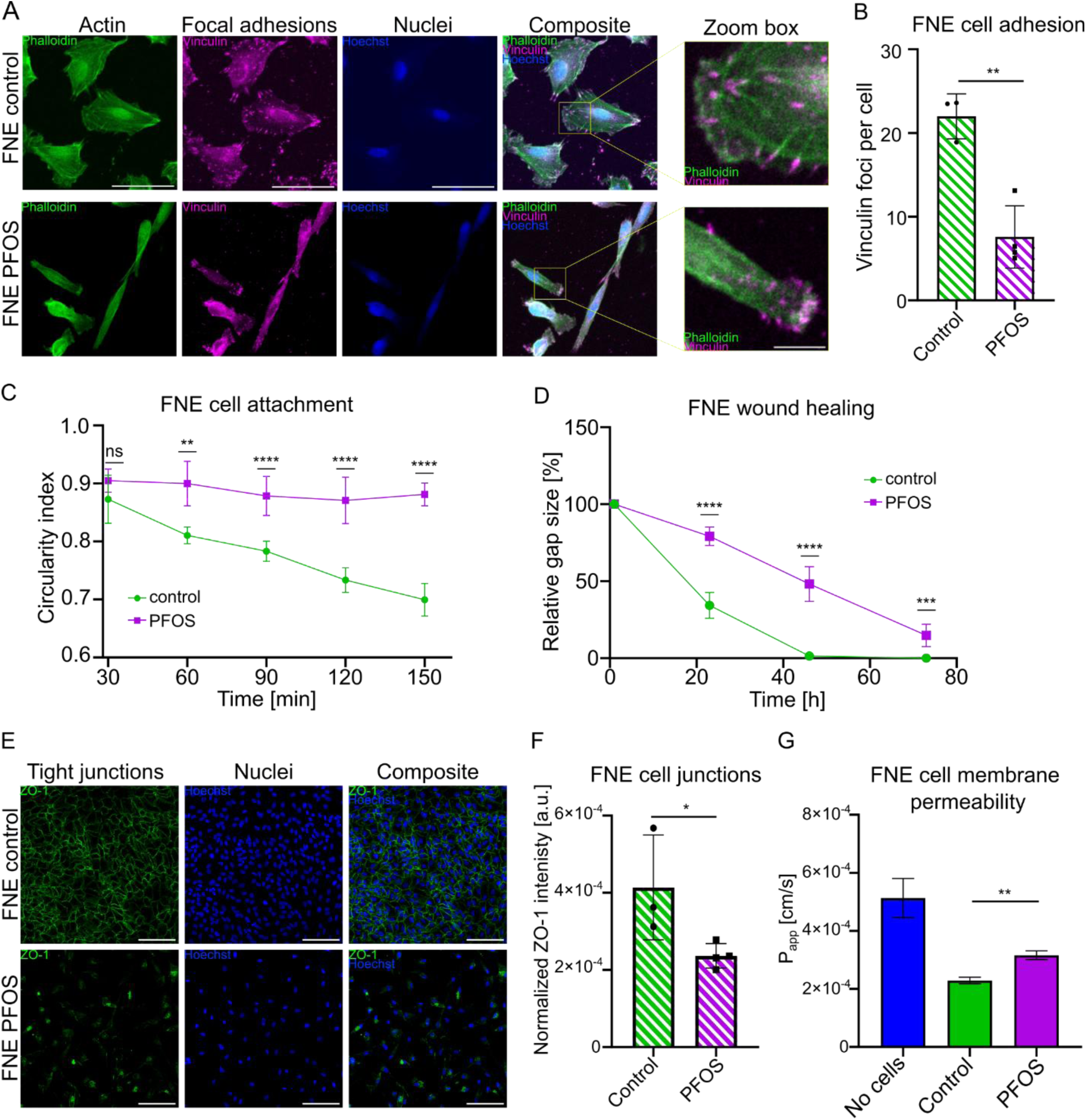
PFOS weakens focal adhesion organization and epithelial barrier integrity in FNE cells. **(A)** Confocal images of FNE cells immunofluorescently stained for F-actin (phalloidin; green), vinculin (magenta), and nuclei (Hoechst; blue) 48 h after treatment with 25 µM PFOS or vehicle control. Zoomed insets highlight actin fiber structures. Scale bars = 50 µm. **(B)** Quantification of vinculin foci per cell averaged across > 50 cells per replicate (n = 3). **(C)** Time course of FNE cell reattachment following brief detachment and re-plating. Circularity index (mean ± SD) was used as a measure of spreading efficiency. **(D)** Collective migration quantified by wound-healing assay. Confluent monolayers were scratched after 48 h treatment ± PFOS and imaged every 12 h. **(E)** Immunofluorescent staining of tight-junction protein ZO-1 (green) and nuclei (Hoechst; blue) showing disrupted junctional continuity in PFOS-treated cells. Scale bars = 50 µm. **(F)** Normalized ZO-1 edge intensity per cell; each point represents one replicate (n = 3). **(G)** Paracellular permeability (P_app_) measured by TRITC-dextran flux across membrane alone (no cells), or confluent monolayers of FNE control (Control) or PFOS-treated FNE (PFOS). Increased P_app_ indicates loss of barrier integrity. All data represent mean ± SD from 3 replicates. Statistical comparisons used unpaired t-tests (B, F, G) or two-way ANOVA with Sidak’s post hoc test (C, E). *p < 0.05; **p < 0.01; ***p < 0.001; ****p < 0.0001.

### PFOS impairs epithelial barrier integrity of FNE cells

Considering that the FNE cells change their shape and adhesion when exposed to PFOS, we investigated whether PFOS impairs the fundamental epithelial function of FNE cells. Using a wound healing assay, we found that PFOS-treated FNE cells exhibited significantly delayed wound closure, indicative of impaired monolayer regeneration (**Figure 2D, S3B**). Beyond slower wound healing, PFOS-treated FNE cells appeared to be disordered and often breaking away from the monolayer, which rarely occurred in vehicle-control cells, where the entire monolayer closed the gap with high level of directionality and coherence (**Figure S3B**; **Movie 3**,**4**). Indeed, immunofluorescent staining revealed reduced ZO-1 intensity at cell edges, suggesting a compromised integrity of epithelial junctions in PFOS-treated FNE cells (**Figure 2E,F**). Because PFOS-treated cells largely failed to establish the typical cobblestone-like epithelial monolayer, the organization of tight junction staining may in part reflect altered cell morphology rather than junctional complex integrity alone. Consistent with the compromised epithelial integrity, we found that paracellular flux, which reports movement of a reporter molecule across an epithelial cell layer between adjacent cells and reflects the permeability of tight junctions, was increased in PFOS-treated FNE cell monolayers (**Figure 2G**, **S3C**). Taken together, the delay in wound healing, weakening of epithelial monolayer integrity, and increase in monolayer permeability are consistent with PFOS-induced epithelial barrier impairment of FNE cells.

### PFOS induces transcriptional dysregulation of genes involved in lipid metabolism, cell adhesion, and signal transduction

To investigate the gene expression changes underlying the observed phenotypes in FNE cells upon PFOS exposure, we performed transcriptomic profiling. RNA sequencing revealed that acute PFOS treatment (25 µM, 48 h) led to a modest but significant transcriptional response, with 82 significantly differentially expressed genes (DEGs) identified (**Figure 3A**). A heat map of the top 20 DEGs highlighted a clear separation between PFOS-treated and control samples, with lipid metabolism–related genes such as APOA1 and SFTPB, cytoskeleton and cell adhesion-related genes such as KRT80 and THBS1, and transcription regulation genes such as ID4, strongly downregulated (**Figure 3B**). Transcripts of ECM modulators such as PSAT1 and ITGB3, stress-response genes such CHAC1 and ASNS, signal transduction-related genes such as DUSP5, and immune-related genes including SERPINB2 and LCN2 were upregulated (**Figure 3B**). While the overall number of DEGs was relatively limited, consistent with the short exposure duration, these genes represented a diverse array of biological processes (**Figure 3C**).

**Figure 3.**
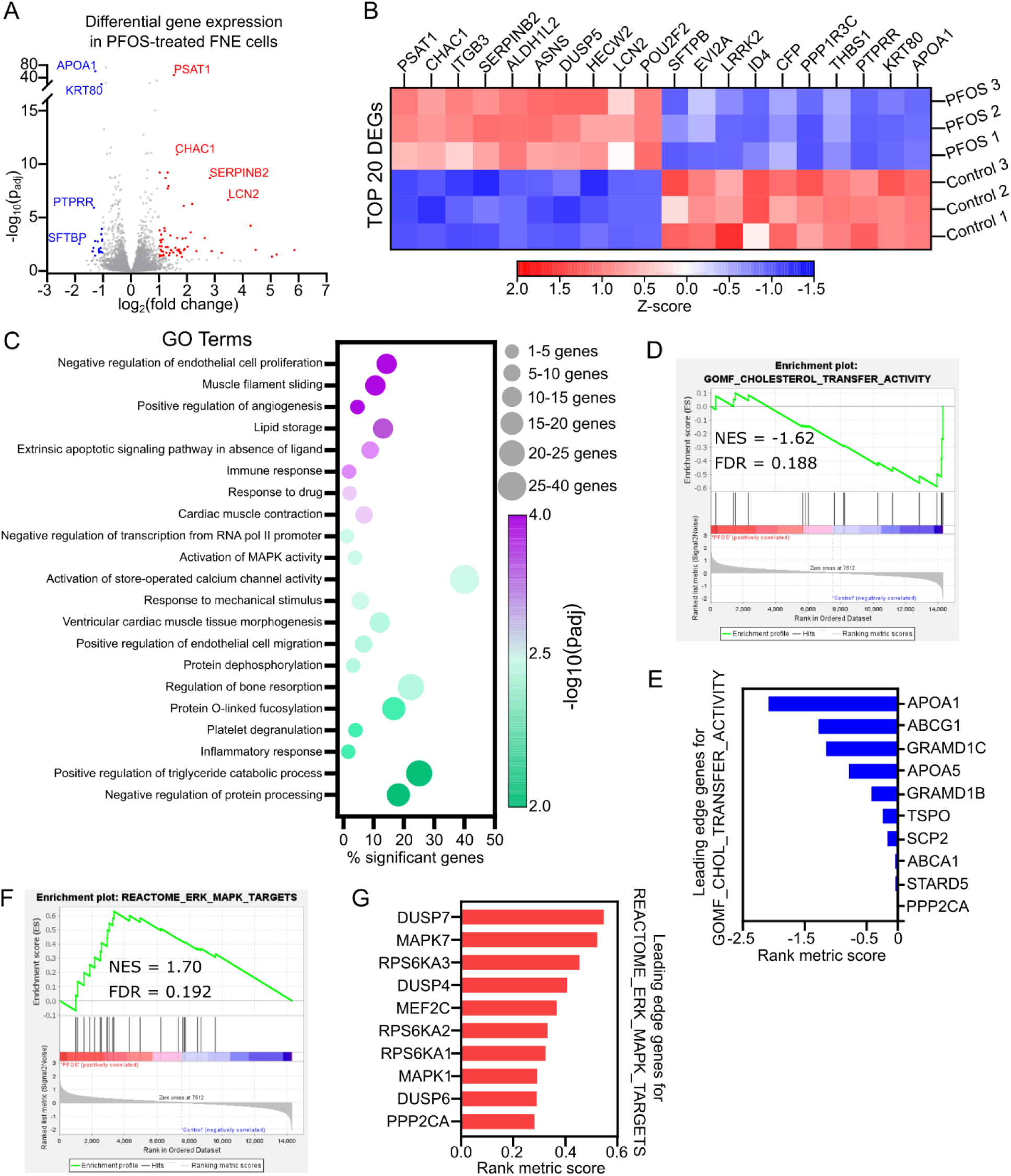
Transcriptomic profiling reveals dysregulation of lipid metabolism, adhesion, and stress-signaling pathways in PFOS-treated FNE cells. **(A)** Volcano plot showing differentially expressed genes (DEGs) in FNE cells after 48 h treatment with 25 µM PFOS versus control. Genes meeting p_adj_ < 0.05 and log_2_(fold change) are highlighted in red (up-regulated) and blue (down-regulated). **(B)** Heatmap of the 20 most significantly altered transcripts (z-score normalized across samples) demonstrating clear segregation between PFOS-treated and control replicates. **(C)** Gene Ontology (GO) enrichment of top 20 most significantly altered biological processes. Bubble size indicates the number of DEGs in each term; color corresponds to –log_10_(p_adj_). **(D)** GSEA enrichment plot for GOMF_Cholesterol_Transfer_Activity, indicating reduced expression of cholesterol-handling genes (NES = –1.62; FDR = 0.188). Negative NES indicates this transcription signature is less enriched in PFOS compared to control FNE cells. **(E)** Leading-edge genes contributing most to the negative cholesterol-transfer pathway enrichment based on rank metric score. **(F)** GSEA plot showing positive enrichment of REACTOME_ERK_MAPK_TARGETS (NES = 1.70; FDR = 0.192). Positive NES indicates this transcription signature is more enriched in PFOS compared to control FNE cells. **(G)** Leading-edge genes contributing to the positive ERK/MAPK target enrichment based on rank metric score.

To interpret the pathways related to these gene expression changes, we conducted Gene Ontology (GO) and Gene Set Enrichment Analysis (GSEA). A summary dot plot of the top enriched GO terms revealed significant clustering of pathways related to lipid storage, regulation of triglyceride metabolism, and protein dephosphorylation, which are often linked to global perturbations in lipid homeostasis, possibly due to membrane changes or disrupted lipid transport (**Figure 3C**). In agreement with our *in vitro* observations, GO pointed toward cytoskeletal remodeling and altered adhesion tension, represented by dysregulated mechanosensitive signaling GO terms such as response to mechanical stimulus, muscle filament sliding, cardiac muscle tissue morphogenesis, and activation of MAPK (**Figure 3C**).

GSEA further demonstrated that acute PFOS exposure attenuated pathways involved in cholesterol transport activity (**Figure 3D**). To further identify key drivers of lipid homeostasis shifts, we examined the core genes that contribute most to the enrichment signal in GSEA. Among these, ***APOA1*** emerged as the top downregulated gene (**Figure 3E**). *APOA1*, a canonical PPARα target gene, encodes apolipoprotein A1, a key component of high-density lipoprotein (HDL) particles, which plays a central role in cholesterol efflux and membrane homeostasis (Mangaraj et al., 2016, Curtiss et al., 2006, Jonas, 2000). The marked downregulation of *APOA1* suggests that PFOS exposure may perturb lipid transport and membrane composition, potentially compromising membrane integrity and triggering compensatory transcriptional responses. Consistent with *APOA1* suppression, GSEA showed suppression of PPARα pathway genes (**Figure S4A**). PFOS has previously been identified as a weak PPARα agonist that selectively induces fatty-acid-oxidation genes while failing to activate LXR/PPARα-dependent apolipoprotein targets in hepatic cells.

In addition, GSEA revealed activation of several membrane-proximal signal transduction pathways. PFOS exposure upregulated ERK-MAPK targets (**Figure 3F,G**), KRAS/MAPK signature (**Figure S4B**) and mTORC1 (**Figure S4C**), while canonical TGF-β/EMT programs were suppressed (**Figure S4D,E**). Several stress-response transcriptional programs that depend on P-TEFb (CDK9) were prominent, including MYC targets, TNFα/NF-κB, E2F/G2M, and the unfolded-protein response (**Figure S4F–J**). Together with broad suppression of ECM adhesion programs (**Figure S4K**), these data indicate that PFOS engages an adhesion–MAPK–transcriptional regulation axis.

Along with the enrichment of E2F and G2/M checkpoint gene sets, we noticed a strong upregulation of DNA damage response, cell-cycle checkpoint signaling and negative regulation of mitotic progression, consistent with the proliferation arrest observed *in vitro* in PFOS-treated FNE cells (**Figure S4L–N**).

### PFOS-induced morphology change is reversed by targeted inhibitors and cholesterol-mediated membrane stabilization

We next tested whether blocking some of the key pathways identified by our transcriptomic profiling could reverse the morphological changes induced by PFOS. Pharmacological inhibition of MEK, largely restored cell circularity and increased cell-substrate adhesion in PFOS-treated FNE cells (**Figure 4A–C, S5A**). In addition to MEK, inhibition of other kinases relevant to pathways identified in our transcriptomic analyses, such as focal adhesion kinase (FAK), cyclin-dependent kinase 9 (CDK9) and Src family kinases (SFKs), showed at least partial rescue of PFOS-induced FNE cell shape alterations (**Figure S5A,C**). These results are consistent with previous data supporting the role of MEK or FAK inhibition in stabilization of focal adhesions (Vomastek et al., 2007, Kanteti et al., 2018). Inhibition of mTORC1/2 on the other hand, did not show a significant rescue, while inhibition of GSK3β further exacerbated the spindly phenotype of PFOS-exposed FNE cells. (**Figure S5B,C**). This indicated that PFOS-induced morphology alterations are mediated, at least in part, by signaling through the adhesion-linked stress-signaling axis, coupling membrane-proximal cues to transcriptional regulation.

**Figure 4.**
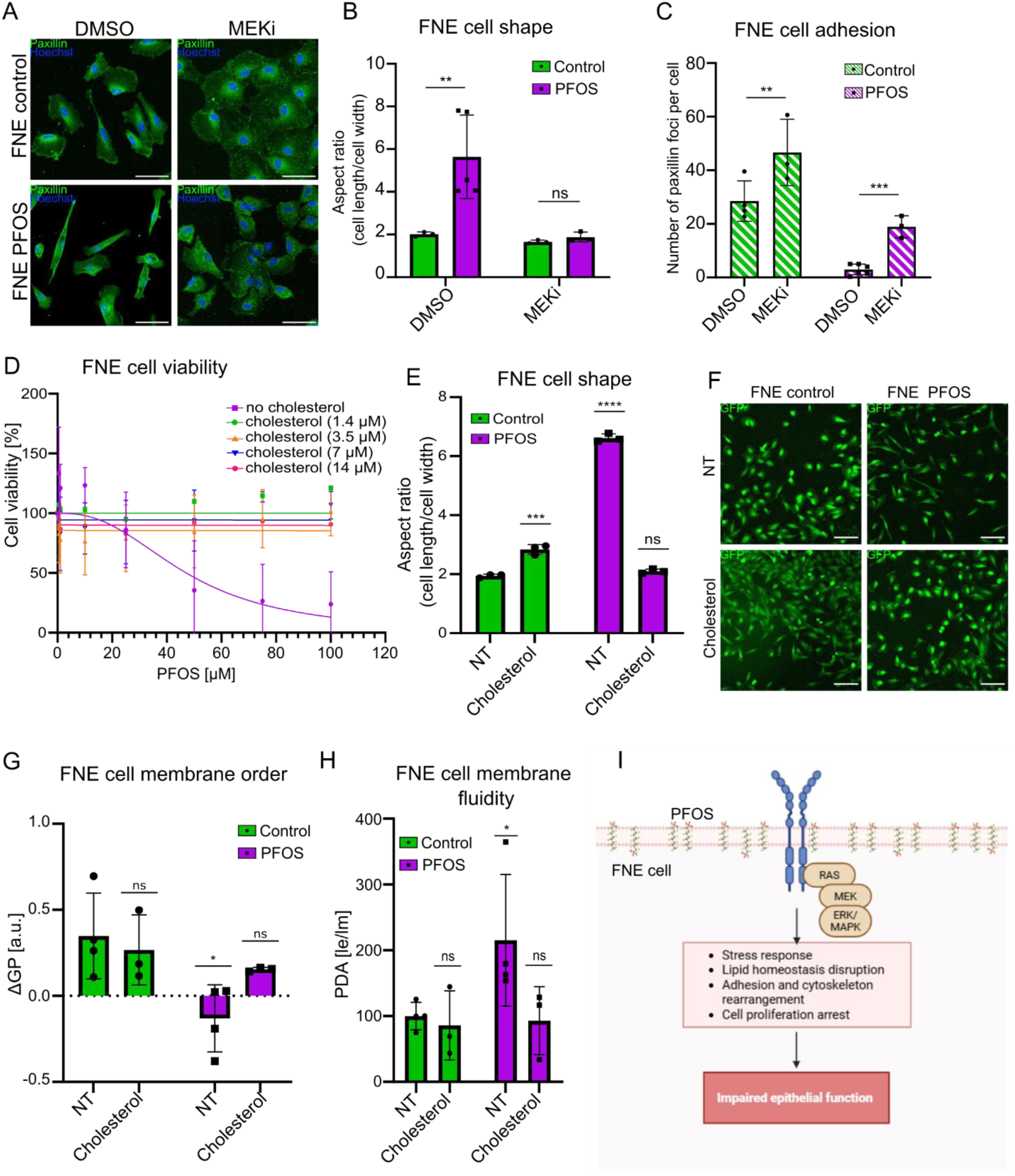
Inhibition of MAPK signaling or cholesterol supplementation rescues PFOS-induced defects. **(A)** Representative fluorescence images of FNE cells stained for focal adhesions (paxillin; green) and nuclei (Hoechst; blue) following 48 h PFOS treatment with or without 24-hour pre-treatment with 0.5 µM MEK inhibitor (PD0325901). DMSO served as vehicle control for MEK inhibitor. Scale bars = 50 µm. **(B)** Quantification of cell aspect ratio (length/width) demonstrating restoration of near-normal morphology by MEK inhibition. **(C)** Quantification of paxillin foci per cell from **(A)** averaged across > 10 cells per replicate (n = 3). **(D)** PFOS cytotoxicity curves ± cholesterol supplementation measured after 96 h by viability assay. Cholesterol was added 24 h before PFOS exposure at the estimated 1.4–14 µM final concentrations (of free cholesterol). **(E)** Quantification of cell aspect ratio in untreated (NT) or cholesterol pre-treated cells ± PFOS showing complete morphological rescue. **(F)** Representative GFP fluorescence images of control and PFOS-treated FNE cells with or without cholesterol pretreatment, illustrating the rescue of spindle-like morphology. GFP signal is from constitutively expressed transgene in FNE cells. Scale bars = 100 µm. **(G)** Membrane order quantified by Laurdan generalized polarization (ΔGP). ΔGP for human cells falls roughly between -1 and +1. Higher ΔGP indicates a more ordered (less fluid) membrane. **(H)** Membrane fluidity determined by PDA excimer-over-monomer (le/lm) fluorescence ratio (higher le/lm ratio = more fluid). **(I)** Shematic model of PFOS disrupting FNE cell membrane, leading to activated stress response, disrupted lipid homeostasis, adhesion and cytoskeleton remodeling, and ultimately resulting in impairment of FNE cell epithelial function. Data are mean ± SD (n = 3–4 biological replicates). Statistical tests: one-way ANOVA with Sidak’s post hoc correction. *p < 0.05; **p < 0.01; ***p < 0.001; ****p < 0.0001.

In contrast, pharmacological modulation of PPARα signaling failed to rescue the PFOS-induced phenotype. Treatment with PPARα antagonists (GW-6471, GW-9662) or agonists (WY-14643, tesaglitazar) did not prevent PFOS-treated FNE cells from adopting the characteristic spindle-like morphology (**Figure S5C,D**). These results indicate that the morphological effects of PFOS are not primarily mediated through PPARα activation, at least not directly or exclusively. This is consistent with prior *in vivo* observations that PPARα knockout only partially mitigates PFAS-induced hepatic toxicity, suggesting that PPARα activation alone cannot account for the broader cellular effects of PFOS, which likely involve membrane perturbation and secondary signaling responses.

Given the propensity of PFOS to perturb membranes, and that cell adhesion signaling and lipid organization control receptor partitioning, our transcriptomic profiling is most consistent with the hypothesis that indirect, membrane-initiated changes could drive signal transduction pathways, rather than PFOS directly affecting dysregulation of the kinase transcripts themselves. We reasoned that supplementing FNE cells with exogenous cholesterol could reinforce their cell membrane making them less susceptible to the potential membrane-disrupting effects of PFOS.

Supplementing even low amounts of cholesterol prior to PFOS exposure completely abrogated the cytotoxic effects of PFOS on FNE cells and restored their shape (**Figure 4D–F**). This was consistent with the idea that PFOS incorporation into the plasma membrane might alter its biophysical properties, triggering aberrant activation of membrane-associated signaling complexes through indirect mechanisms such as receptor clustering, lipid raft disruption, or altered lateral diffusion (Lingwood and Simons, 2010, Kusumi et al., 2011).

To quantitatively test changes in the properties of the plasma membrane, we assessed membrane order by Laurdan dye generalized polarization (GP) (Harris et al., 2002, Levitan, 2021) which revealed that PFOS-treated FNE cells exhibit a more disordered membrane organization compared to control FNE cells (**Figure 4G**). Consistent with this, an independent assay using a <u>p</u>yrene<u>d</u>ecanoic <u>a</u>cid (PDA)-based membrane probe demonstrated increased membrane fluidity following PFOS exposure (**Figure 4H**). Notably, even a short cholesterol pulse administered after PFOS exposure partially restored membrane order and fluidity. (**Figure 4G**, **4H**). These findings strongly suggest that PFOS perturbs FNE cells membrane organization and disrupts lipid homeostasis, leading to activation of downstream signaling pathways that mediate stress response and ultimately results in alterations in cell adhesion and cytoskeleton, impairing their epithelial functions (**Figure 4I**).

## DISCUSSION

Our findings demonstrate that acute PFOS exposure disrupts key aspects of fallopian tube epithelial cell biology, including cell morphology, proliferation, motility, and epithelial barrier integrity. These effects were accompanied by transcriptomic dysregulation across pathways, from elevated stress response to reduced cholesterol transport. Pharmacological inhibition of MEK signaling rescued PFOS-induced shape changes and FNE cell substrate adhesion. Furthermore, exogenous cholesterol supplementation reversed the phenotype associated with PFOS exposure, suggesting that PFOS integration into plasma membrane could disrupt membrane-proximal signaling, including the KRAS/MEK/ERK pathway.

Prior work shows that many amphiphilic PFAS compounds readily partition into lipid bilayers and can incorporate into membranes where they alter packing, permeability, and order, both in model systems and in cells (Hu et al., 2003, Zhao et al., 2023). Consistent with a membrane-mediated mechanism, we observed decreased membrane order and increased membrane fluidity in FNE cells following PFOS exposure. Exogenous supplementation of FNE cells with cholesterol, which is known to increase lipid packing, order and generally stiffen phospholipid bilayers, mitigated PFOS-induced shape changes and cytotoxicity (Chakraborty et al., 2020, Henriksen et al., 2006). Brief cholesterol supplementation following PFOS exposure could reverse membrane fluidity and disorder, suggesting that direct modulation of membrane composition can restore biophysical properties. Together, these results support a model in which PFOS perturbs membrane fluidity thereby likely destabilizing membrane-associated signaling complexes or promoting non-specific receptor clustering, and ultimately leading to aberrant activation of downstream signaling pathways (**Figure 4I**).

Membrane perturbation as an underlying mechanism of PFAS toxicity is increasingly recognized, though it remains less emphasized than mechanisms focusing on protein targets (Bangma et al., 2022). Our study extends this concept to human fallopian tube epithelium and links it to focal adhesion signaling, reattachment kinetics, and epithelial dysfunction. Given the fallopian tube’s role in reproduction and disease susceptibility, defining how membrane composition reacts to PFOS exposure could have practical value. Membrane-focused interventions (e.g., controlled sterol supplementation or lipid-environment modulation) may provide mechanistic insights and suggest therapeutic strategies to blunt acute PFOS injury in a broad variety of epithelial tissues.

## METHODS

### Cell culture

Unless otherwise indicated, FNE cells (wild-type and ΔTP53) were maintained in a mixture of 42% Medium 199, 21% DMEM and 21% F-12 (2:1:1 mixture) basal medium and supplemented with 1% penicillin/streptomycin antibiotic, 2% FBS, 14 µg/mL insulin, 27 ng/mL cholera toxin (holotoxin), 560 ng/mL hydrocortisone, 220 ng/mL T3, 55 ng/mL β-estradiol, 83 ng/mL all-trans retinoic acid (ATRA), and 11 ng/mL epidermal growth factor (EGF). RPE1 cells and ovarian carcinoma cell lines (CaOV3, Kuramochi, RMUGS) were cultured in 48.5% DMEM and 48.5% F-12 (1:1 mixture) basal medium, supplemented with 1% penicillin/streptomycin antibiotic and 2% FBS for cross-assay comparability. Where indicated, 0–10% FBS titrations in OptiMEM (Gibco) as basal medium were used to assess serum effects. All cells were grown at 37 °C in 5% CO₂ conditions.

### PFOS handling

PFOS (perfluorooctane sulfonate; Santa Cruz) was prepared as a 5 mM stock in sterile, molecular biology-grade water using polystyrene tubes both for storage, dilution, and mixing to minimize adsorption. PFOS stock solutions in water were stored at 4°C and used for no longer than 4 weeks. Working solutions were freshly diluted in pre-warmed medium; final concentration of vehicle (water, DMSO) in medium was ≤5% for water and ≤0.5% for DMSO. Because protein binding reduces free PFOS, experiments were performed in 2% FBS (unless otherwise noted).

### Cell treatment

For acute exposure, cells received a single dose of 25 µM PFOS for 48–96 h. For washout, cultures were rinsed 3× with warm 1X phosphate buffered saline (PBS) and returned to PFOS-free medium for >24 h. Where indicated, cells were pre-coated on substrates: fibronectin 1–100 µg/mL (Sigma-Aldrich, F1141-1MG), laminin 100 µg/mL (Corning, 354239), or poly-L-lysine 0.01% (Sigma-Aldrich) for ≥1 h at 37 °C prior to seeding. Pharmacologic inhibitors were used at the indicated final concentration:

| Reagent | Target / modality | Stock solution concentration | Vendor / cat. # | Final concentration used |
| --- | --- | --- | --- | --- |
| **PF-573228** | p-FAK / inhibitor | 100 mM | ApexBio<br>B1523 | 10 μM |
| **GSK2256098** | p-FAK / inhibitor | 10 mM | Cayman<br>Chemical<br>Company<br>22955 | 5 μM |
| **CHIR99021** | GSK3β / inhibitor | 10 mM | Cayman<br>Chemical<br>Company<br>13122 | 10 μM |
| **GW 6471** | PPARα /<br>antagonist | 10 mM | Cayman<br>Chemical<br>Company<br>11697 | 10 μM |
| **GW 9662** | PPARγ /<br>antagonist | 10 mM | Cayman<br>Chemical<br>Company<br>70785 | 10 |
| **Wy 14643** | PPARα agonist | 100 mM | Cayman<br>Chemical<br>Company<br>70730 | 10 μM |
| **Teasglitazar** | dual PPARα & PPARγ / agonist | 10 mM | Cayman Chemical Company<br>16791 | 5 μM |
| **Rapamycin** | mTORC1 / inhibitor | 10 mM | Ambeed Inc.<br>A656002 | 0.1 μM |
| **PD0325901** | MAPK1 (MEK/ERK2) / inhibitor | 1 mM | MedChem Express<br>HY-10254 | 0.5 μM |
| **Saracatinib** | Src / inhibitor | 10 mM | Selleck Chemicals<br>379231-04-6 | 5 μM |
| **Dinaciclib** | CDK9 / inhibitor | 37.831 mM | Selleck Chemicals<br>S2768 | 0.1 μM |

All inhibitors or compounds were added 24 hours prior PFOS exposure. Vehicle controls matched DMSO content, which was kept at ≤0.5%. To increase membrane rigidity and decrease sensitivity to PFOS, cells were pre-treated (24 hours prior PFOS exposure) with cholesterol in form of a cholesterol lipid concentrate (Thermo Fisher, 12531018), diluted in complete medium to low working doses (0.05-1X of manufacturer’s recommended final dilution corresponding to an estimated 1.4-28 µg/mL of free cholesterol). Rescue of morphology and viability was assessed after 48–96 h. For acute membrane effects on membrane order and fluidity, a pre-mixed cholesterol–MβCD complex (5 mM cholesterol with 75 mM MβCD or 1:15 molar ratio) in HEPES buffered saline (HBS) solution replaced PFOS-containing medium (48 hours post PFOS exposure) and incubated for 30 minutes at 37 °C in 5% CO₂ conditions. Membrane order and fluidity were then assessed as described using Laurdan dye (Invitrogen) or pyrenedecanoic acid (PDA)-based membrane fluidity kit (Abcam).

### Cell viability assays

Dose–response curves were generated by exposing cells (96-well plates, 2-3 technical replicates per dose) to PFOS for 72 h followed by incubation with Alamar Blue for ≤ 4 h, as per manufacturer’s instructions. Fluorescence was read on a Molecular Devices i3x spectrophotometer/fluorometer. IC₅₀ values were calculated with non-linear fit with a four-parameter variable model (GraphPad Prism), excluding outliers by prespecified criteria.

### Flow cytometry

We performed flow cytometry to estimate suspended cell volume of PFOS-treated or vehicle-control cells. Cells were detached with 0.05% trypsin, collected by centrifugation, washed with 1X PBS, and resuspended in 1X PBS containing 2% FBS. Cells in suspension were analyzed on Attune NxT Flow Cytometer (Thermo Fisher) using FSC-A as a size proxy. Unstained size calibration beads (1-15 µm; Thermo Fisher) were run using the same voltage and gain settings. A minimum of 1000 events per sample were acquired after doublet discrimination (FSC-A vs FSC-H) and analyzed with FlowJo v10.10.0 (FlowJo, Ashland, CA, USA). A standard curve based on the size calibration beads was then used to convert sample FSC-A measurements to theoretical cell diameter using linear regression. Cell volume was then derived from the extrapolated cell diameter, using the equation below:

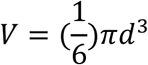

under the assumption the cells take a perfect spherical form in suspension.

### Mass spectrometry

Cell pellets were extracted in methanol, cleared at 16,000 g (10 min), and supernatants dried under N₂. Intracellular PFOS was quantified using an Agilent 1100 HPLC system coupled to a Micromass Quattro Ultima quadrupole mass spectrometer, controlled by MassLynx V4.0 software. The column oven temperature was maintained at 40 °C. An Xterra MS C18 analytical column (150 mm × 4.6 mm; 5.0-μm particle size, from Waters) and Phenomenex delay column (50 mm × 4.6 mm; 3.0-μm particle size, from Phenomenex) were used. The mobile phase consisted of aqueous ammonium acetate (20 mM) (solvent A) and acetonitrile (solvent B) and was pumped at a flow rate of 1.0 mL/min. The starting condition (10% B) was kept for 1 min. The proportion of B was increased linearly to 80% in 5.5 min and kept for 0.5 min. Subsequently, the mobile phase was adjusted to its initial composition in 1 min and held for 3.5 min for re-equilibration, resulting in a total run time of 11.5 min. Fifty microliters of each sample was injected on the column. The MS instrument was operated in the ESI negative mode and the data were acquired in multiple reaction monitoring (MRM) mode. The MS tune parameters and compound parameters were optimized and determined with a syringe pump at a flow rate of 20 µL/min. The capillary voltage was 3.00 kV. The source temperature was 120 °C and the desolvation temperature was 300 °C. The cone voltage was 80.00 V for PFOS. A collision energy of was 10.00 eV for used for PFOS analysis and quantification.

### Epithelial barrier permeability assay

Paracellular flux was measured using 24 mm Transwell® inserts with 0.4 µm pore polycarbonate membrane (Corning, 3412). Polycarbonate membrane material was selected to minimize PFOS adsorption. FNE cells were seeded to confluence and then treated ± PFOS for 48 hours, as described above. Medium ± PFOS was then removed and TRITC-dextran (4 kDa; 0.25 mg/mL; MedChem Express) in Hank’s balanced salt solution (HBSS; Corning) was added to the apical chamber. Basolateral medium was replaced with HBSS alone and sampled at 0–360 min at 15, 30, or 60 minute interval. Fluorescence of TRITC-dextran in basolateral solution was measured (Excitation 485 nm, Emission 520 nm). Apparent permeability (P_app_) was calculated as:

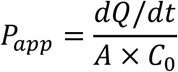

where dQ/dt is the linear flux, A is membrane area, and C_0_ is initial dextran concentration (in apical chamber).

### Immunofluorescent staining

Cells on glass coverslips or glass-bottom imaging plates were fixed in 4% PFA (10 min) or ice cold 1:1 methanol:acetone mixture (10 minutes), permeabilized with 0.5% Triton X-100 (2 min), and blocked in 3% BSA (45 min). The following primary antibodies and stains were used: ZO-1 (Santa Cruz) paxillin (Thermo Fisher) or vinculin (Millipore) for focal adhesions, and FITC-phalloidin (Sigma) for F-actin. Species-matched secondary antibodies with Alexa Fluor were used at 1:2000 dilution (Thermo Fisher). Nuclei were counterstained with Hoechst 33342 (Invitrogen). Samples were preserved in VectaShield antifade mounting medium (Vector Labs), cured overnight, and imaged on confocal microscope.

### Confocal microscopy

Images were acquired on a LSM 880 laser-scanning confocal microscope (Zeiss) with a 20× objective (1.4 NA). Laser lines were used as following: HeN 405 nm (Hoechst), Argon 488 nm (Alexa Fluor 488 or TRITC-phalloidin), Argon 561 nm (Alexa Fluor 568). Pixel size, averaging, and z-step were set using Zen acquisition software. Detector gain/offset were kept constant within experiments. Multiple z-stacked images were collected with the interval of 0.8 µm.

### Live-cell time-lapse microscopy

Still images or time lapses in phase-contrast or fluorescent channels were performed on live cells using BioTek Lionheart FX automated microscope (Agilent), which contains an on-stage incubator (37 °C), with CO₂ (5%) injector. For time lapse microscopy, images were acquired every 20–30 min for 48–96 h using Gen5 image acquisition software (Agilent, version 11.6).

### Quantitative image analysis

All quantitative image analyses were quantified using FIJI (FIJI is just ImageJ) software (Schindelin et al., 2012). Cell length/cell width ratios were derived from Aspect Ratio using best-fit ellipses. FNE cell circularity was assessed by applying consistent segmentation thresholds based on FNE cells constitutively expressing GFP, and measuring circularity index and roundness (FIJI, analyze menu). Cell confluency and cell tracking were analyzed using TrackMate FIJI plugin (Ershov et al., 2022, Tinevez et al., 2017). LoG detector and LAP detector were used to count and track cells (“spots”) over the course of the time lapse. For trajectory analysis, tracks with ≥7 frames were included. Mean speed, mean squared displacement (MSD), and confinement ratio were computed using TrackMate and applied to Trajectory Classifier FIJI plugin (Wagner et al., 2017). Focal adhesions were segmented from paxillin/vinculin channels using background subtraction and thresholding. Only features between > µm²< were selected to focus on focal adhesion foci. Junction continuity was measured as ZO-1 edge intensity, normalized to number of cell nuclei. For wound healing analysis, a manual ROI in phase contrast was used to compute percent closure over time, normalized to the initial ROI size (100%).

### RNA sequencing

Adhered FNE cells were treated with 25 µM PFOS or vehicle (water) for 48 hours. A total of 3 independent samples were prepared per condition. Total RNA was then extracted using the Quick-RNA™ MiniPrep kit (Zymo Research, R1055) following the manufacturer’s protocol (with cell scraping as homogenization method). The RNA concentration and purity were evaluated using a NanoDrop™ 8000 Spectrophotometer (Thermo Fisher Scientific, Fair Lawn, NJ, USA). RNA samples were stored at −80 °C and shipped on dry ice to Genewiz (Azenta US, South Plainfield, NJ, USA) for RNA sequencing. The RNA Integrity Number (RIN) ranged from 8.2 to 9.2 across the samples sequenced. Libraries were prepared according to the company’s standard RNA-seq protocols. Sequencing was performed on an Illumina® HiSeq® platform (Illumina, San Diego, CA, USA; configuration HiSeq 2 × 150 PE HO HiSeq 2 × 150 bp).

### Transcriptomic analysis

The raw reads were assessed for quality and trimmed using Trimmomatic v0.36 to remove adapter sequences and low-quality bases. The clean reads were then aligned to the *Homo sapiens* GRCh38 reference genome (ENSEMBL) using the STAR aligner v2.5.2b, generating alignment files in BAM format. These BAM files served as the basis for quantifying gene expression and performing subsequent differential expression analysis.

Raw sequence reads were assessed for quality and trimmed to remove adapter sequences and nucleotides with poor quality using Trimmomatic v.0.36. Trimmed reads were mapped to the *Homo sapiens* GRCh38 with ERCC genes reference genome available on ENSEMBL using the STAR aligner v.2.5.2b. BAM files aligning the entire read sequences were generated. Unique gene hit counts were calculated using featureCounts from the Subread package v.1.5.2. The hit counts were summarized and reported using the gene_id feature in the annotation file. Only unique reads that fell within exon regions were counted. After extraction of gene hit counts, the gene hit counts table was used for downstream differential expression analysis. Using DESeq2, a comparison of gene expression between the customer-defined groups of samples was performed. The Wald test was used to generate p-values and log2 fold changes. Genes with an adjusted p-value < 0.05 and absolute log2 fold change > 1 were called as differentially expressed genes for each comparison. Gene ontology analysis was performed on the statistically significant set of genes by implementing the software GeneSCF v.1.1-p2. The goa_human GO list was used to cluster the set of genes based on their biological processes and determine their statistical significance. A list of genes clustered based on their gene ontologies was generated. Gene set enrichment analysis (GSEA) was performed using GSEA_4.2.2 software (Subramanian et al., 2005) against selected gene sets from the Molecular Signatures Database (MSigDB 7.5) (Liberzon et al., 2015). The Signal2Noise metric was applied for ranking, and 1000 permutations were used. Results with a false discovery rate (FDR) < 0.25 were considered significant.

### Membrane order measurements

Laurdan generalized polarization (GP) was used to quantify membrane order. Laurdan (Invitrogen) stock solution was prepared fresh at 2 mM in dimethylformamide (DMF). Cells were incubated with 5 µM Laurdan in HBSS for 30 min at 37 °C, washed, and measured on a plate reader (Ex 350–360 nm; Em 440 ± 10 nm and 500 ± 10 nm). ΔGP was calculated as:

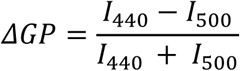

with background subtraction from unstained control cells. Values were averaged per biological replicate and expressed relative to the vehicle mean.

### Membrane fluidity measurements

Membrane fluidity was assessed with a polarity-sensitive PDA-based Membrane fluidity kit (Abcam, ab189819), following the manufacturer’s protocol with optimization for FNE cells. Cells were labeled in perfusion buffer containing 5 µM fluorescent reagent and 0.08% Pluronic F-127 for 1 hour at room temperature. After a gentle wash with HBSS, plates were read from the bottom in black-walled, clear-bottom 96-well plates using a Molecular Devices i3x spectrophotometer/fluorimeter (Ex 380 nm, Em 470 nm for the monomer (lm; ordered emission) and 500 nm for the excimer (le; disordered emission)). Gain was fixed across plates. After background subtraction, fluidity index was computed as the ratio of excimer (le)/monomer (lm) channels (higher ratio = greater excimer formation thus greater fluidity) and normalized to the vehicle mean per plate.

### Statistical analysis

All data are represented as mean ± standard deviation (SD) unless stated otherwise. Number of experimental repeats refers to biological replicates (independent cultures); any technical replicates were averaged within each experimental repeat. Two-group comparisons used unpaired, two-tailed, parametric t-tests and adjustment for multiple comparisons (Benjamini–Hochberg method). Multi-group comparisons were assessed using multiple t-tests or one-way ANOVA with Tukey post hoc tests. Effect sizes and exact p-values are reported in figures or figure legends. Curve fitting (IC₅₀) used a four-parameter variable slope model. All statistical analysis was performed using GraphPad Prism v9.5.1 (GraphPad Software, San Diego, CA, USA).

## Acknowledgements

We would like to thank Dr. Eustace Fernando (former Postdoctoral Fellow, Sarkar Lab) for his assistance with the mass spectrometry analysis. We acknowledge the Department of Chemistry and Chemical Biology at Stevens Institute of Technology for access to instrumentation and technical support.

