## Supplementary Data for "PFOS Disrupts Membrane Signaling and Epithelial Integrity in Fallopian Tube Cells"

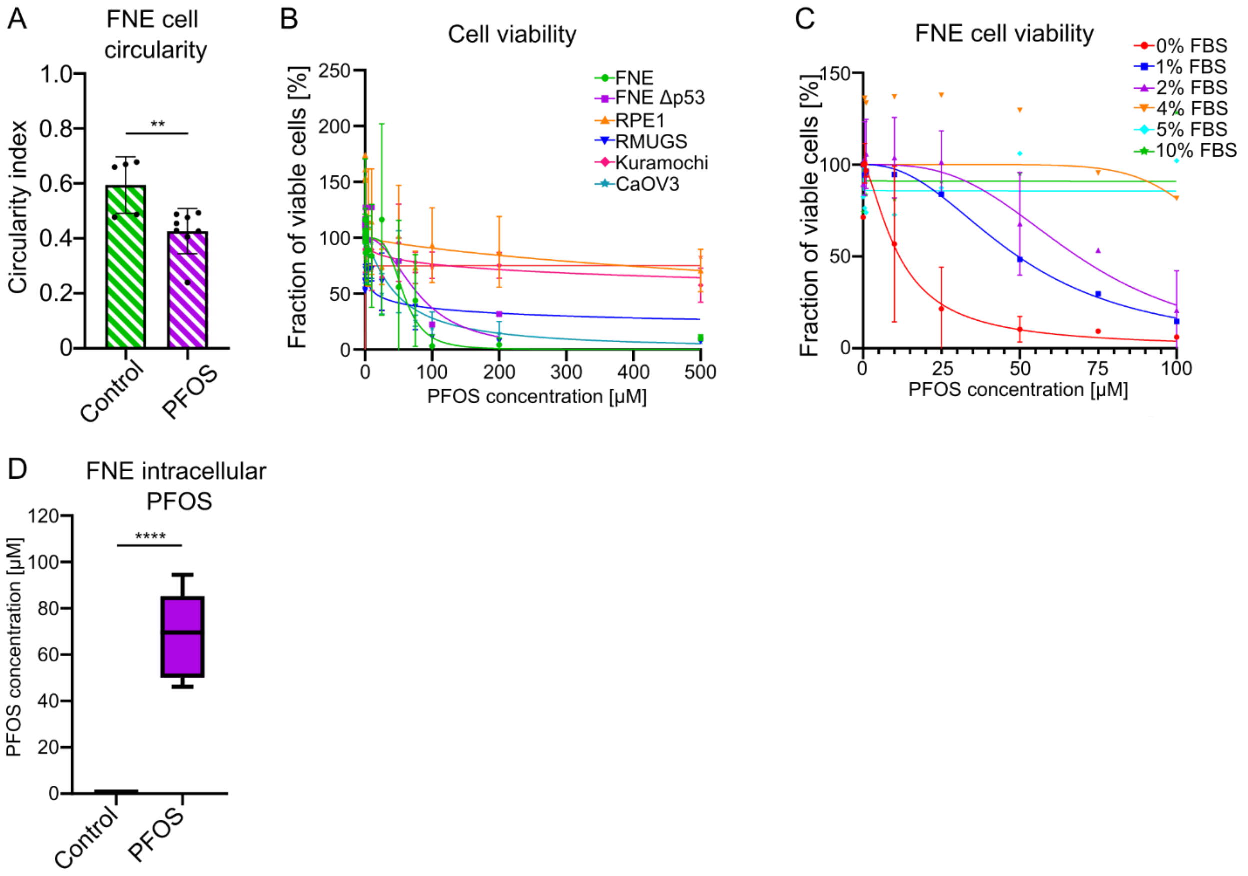


**Figure S1. PFOS induces cytotoxicity in epithelial cells.** (**A**) Circularity index of GFP-expressing FNE cells measured 48 h after PFOS exposure. Each dot represents the mean of > 50 cells per replicate (n = 3). **(B)** Dose-response curves for PFOS cytotoxicity (72 h) in FNE, FNE ΔTP53, RPE1, RMUG-S, Kuramochi, and CaOV3 cells. (C) PFOS $IC_{50}$ values determined under varying fetal-bovine-serum (FBS) concentrations (0–10%) demonstrating FBS-dependent attenuation of PFOS efficacy. **(D)** Intracellular PFOS levels quantified by LC–MS/MS after 48 h exposure to 25 µM PFOS, showing ~300-fold lower accumulation (≈ 80 nM) than extracellular concentration. All data represent mean ± SD (n = 3 independent repeats). *$p < 0.05$; **$p < 0.01$; ****$p < 0.0001$.

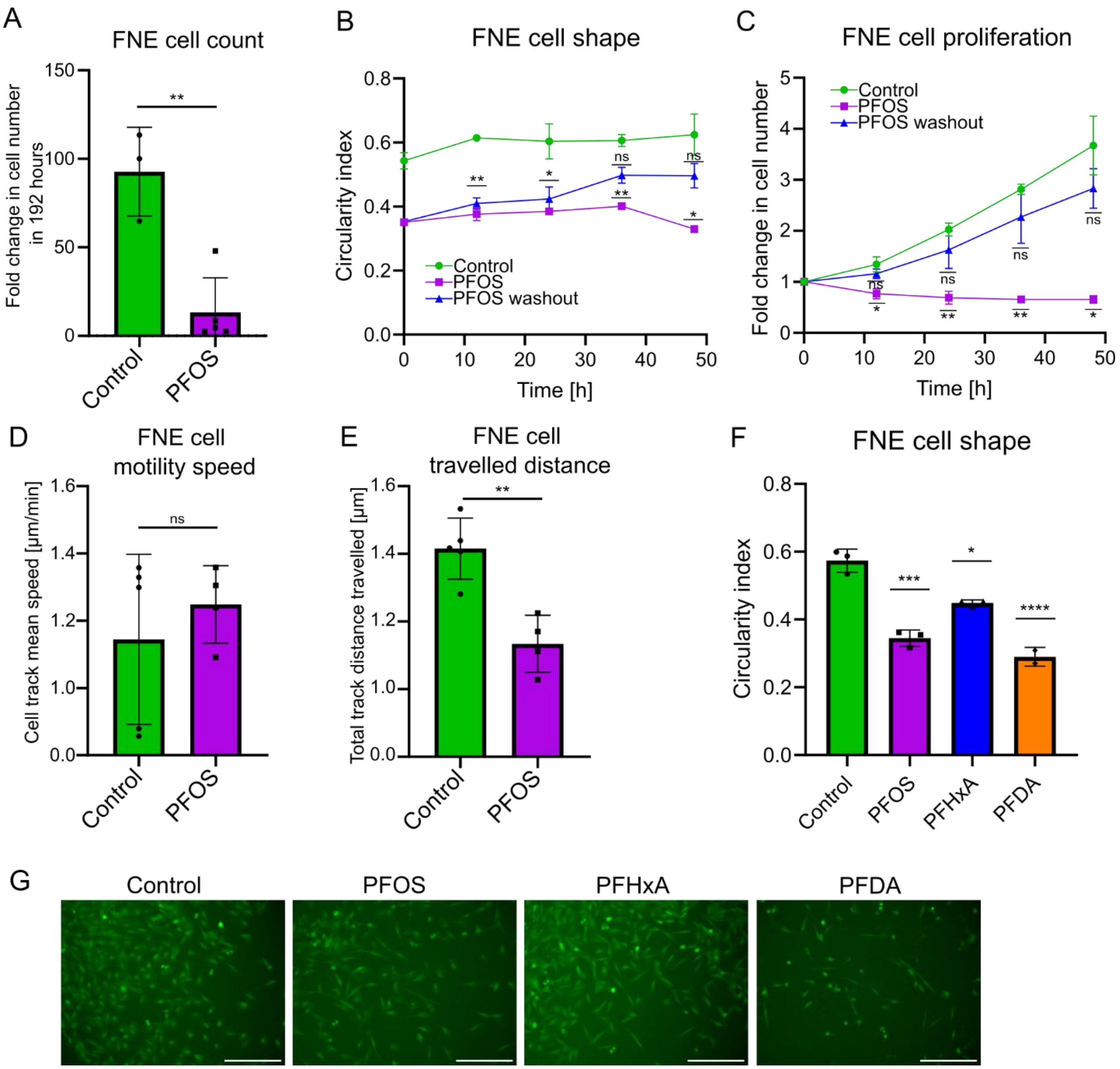


**Figure S2. PFOS induces reversible morphological and motility changes in FNE cells.** (**A**) Fold change in total FNE cell number after 192 h ± PFOS. **(B)** Time-lapse quantification of FNE morphology (circularity index) and **(C)** proliferation following PFOS washout, showing rapid restoration of growth and cell shape within 24-48 h of PFOS removal. Each time point represents the mean of > 10 cells per replicate (n = 2). **(D)** Mean speed of individual FNE cell tracks over time extracted from live-cell imaging of GFP-expressing cells ± PFOS. **(E)** Quantification of average net displacement (end-to-end distance or total distance travelled) per sample over 24 h. Each dot represents an average of all cell tracks in the sample (≥ 50 tracks per replicate, n = 3). **(F)** Circularity index of GFP-expressing FNE cells measured 48 h after exposure to vehicle control (water + DMSO) or various PFAS: PFOS, PFHxA, or PFDA. Each dot represents the mean of > 10 cells per replicate (n = 3). **(G)** Representative fluorescent images of FNE cells constitutively expressing GFP that were exposed to various PFAS (PFOS, PFHxA, or PFDA) or vehicle control (water + DMSO). Cells were treated with PFAS or vehicle-control for 48 hours. All data represent mean ± SD. *$p < 0.05$; **$p < 0.01$; ****$p < 0.0001$.

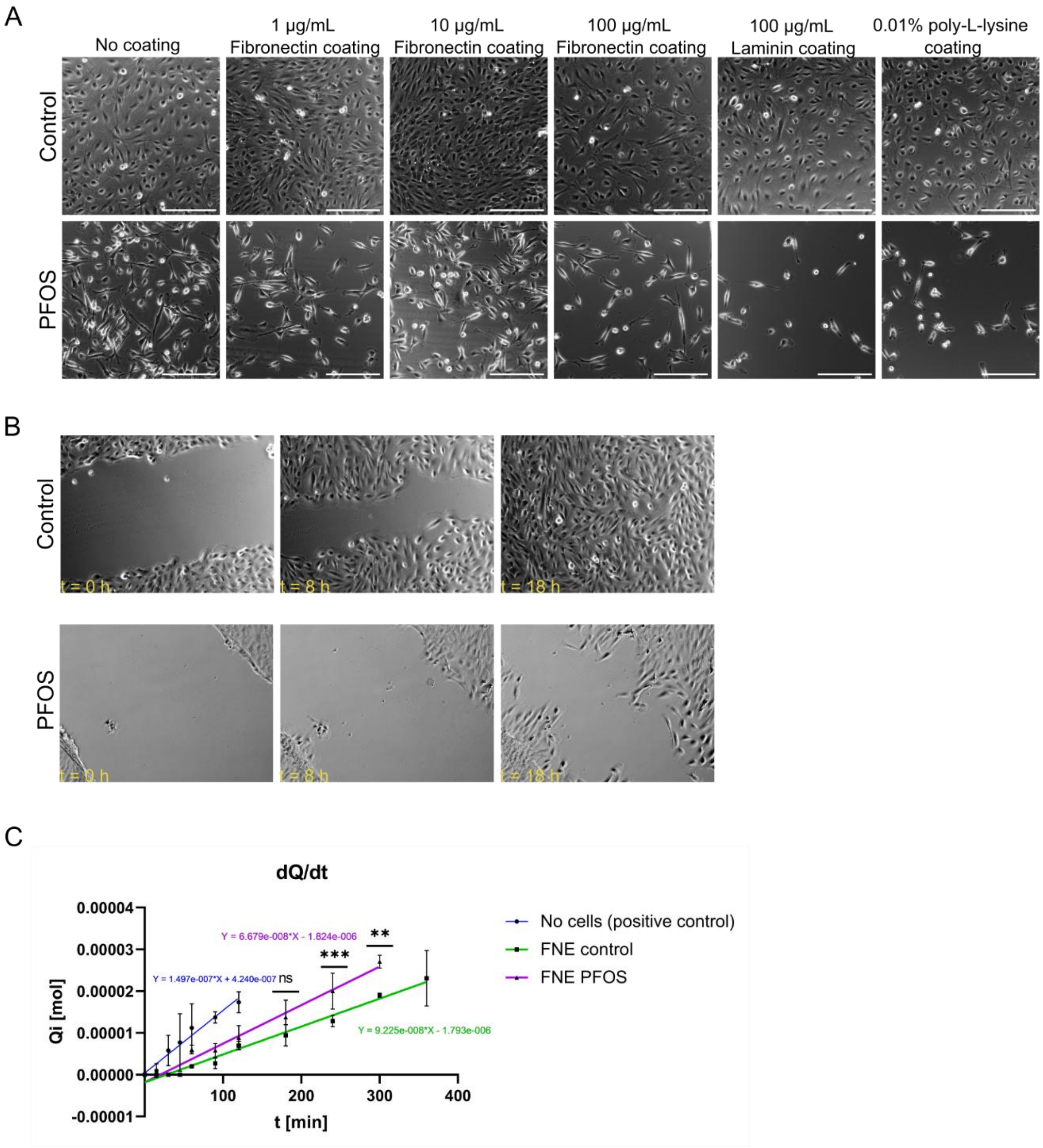


**Figure S3. PFOS weakens focal adhesion organization and epithelial barrier integrity in FNE cells. (A)** Representative phase contrast images of FNE cells grown on pre-coated surfaces (with 1X PBS (negative control), 1 µM Fibronectin (FN), 10 µM Fibronectin (FN), 100 µM Fibronectin (FN), 100 µM Laminin, or 0.01% (w/v) of poly-L-lysine) and exposed to PFOS or vehicle control (water). FNE cells were treated with PFOS or vehicle-control for 48 hours. Scale bar = 100 μm. **(B)** Representative phase contrast images of the wound healing assay. FNE cells were grown to form a monolayer, then exposed to PFOS or vehicle control (water) for 48 hours, scratched, and imaged every 20 minutes for 24 hours. **(C)** Time-resolved paracellular flux of TRITC–dextran (4 kDa) across confluent FNE monolayers ± PFOS (green and magenta) or membrane insert alone (blue). Fluorescence in the basolateral chamber (Q) was measured at 15–60 min intervals over 6 h and plotted as cumulative dextran passage versus time. Linear regression of the linear portion of the curve (0–120 min for membrane alone, 0–300 min for FNE + PFOS, and 0–360 min for FNE control) was used to determine the flux rate (dQ/dt) for each condition, which was then used to calculate apparent permeability (Papp) shown in **Figure 2G**. PFOS-treated monolayers exhibited a steeper slope, indicating higher dextran flux and compromised barrier integrity. All data represent mean ± SD (n = 3). *p < 0.05; **p < 0.01; ****p < 0.0001.

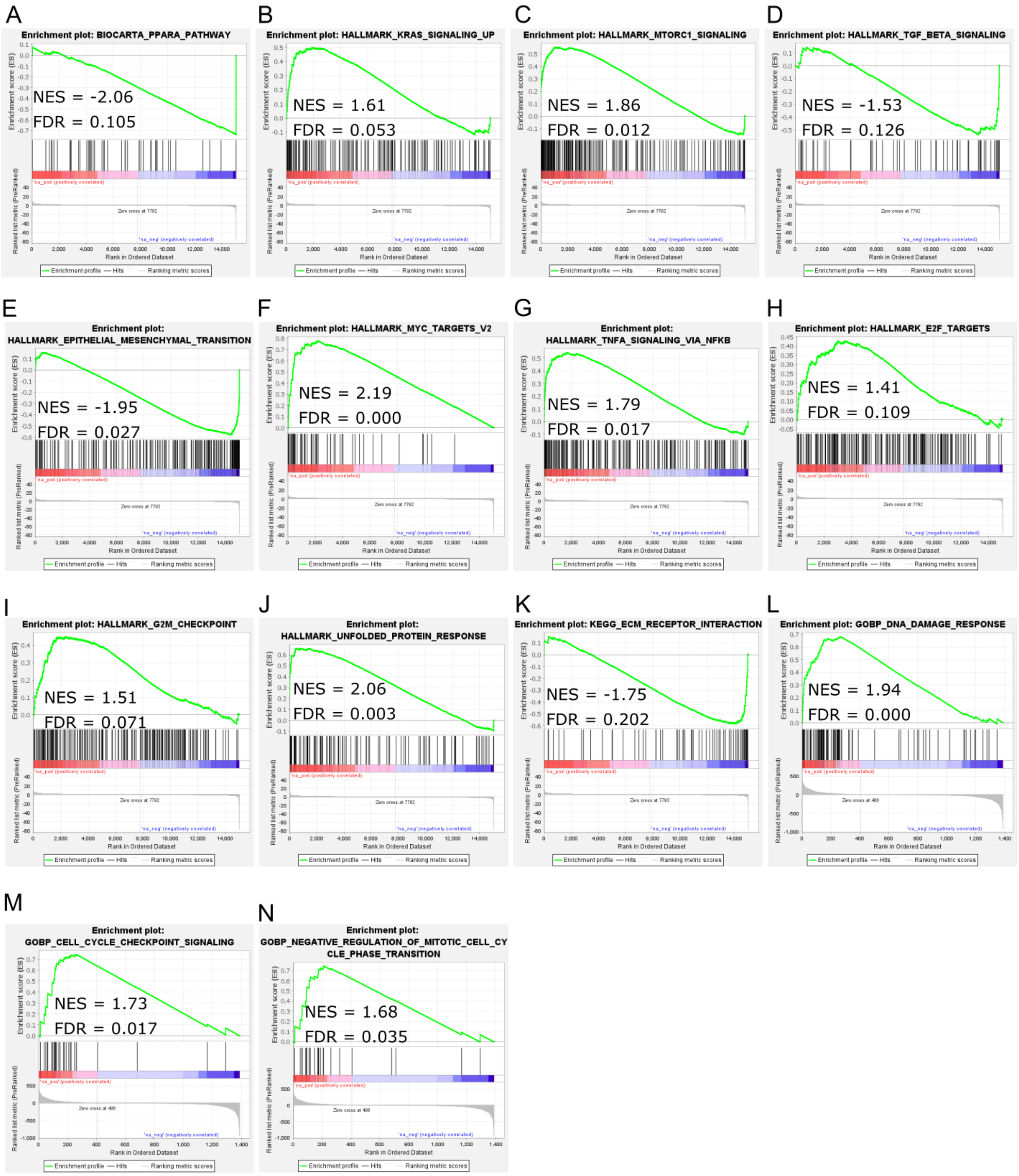


**Figure S4. Gene set enrichment analysis identifies activation of MAPK, MYC, and stress-response pathways in PFOS-treated FNE cells. (A–N)** Representative GSEA enrichment plots showing significantly altered transcriptional programs in PFOS-treated versus control FNE cells. Positively enriched gene sets in PFOS-treated FNE cells (indicated by positive NES) included

KRAS/MAPK, mTORC1, MYC targets, TNFα/NF-κB, unfolded-protein response, G2/M checkpoint, cell-cycle checkpoint signaling, and DNA-damage response, whereas PPARα and EMT programs were negatively enriched in PFOS-treated FNE cells (indicated by negative NES). Normalized enrichment scores (NES) and false discovery rates (FDR) are indicated on each plot.

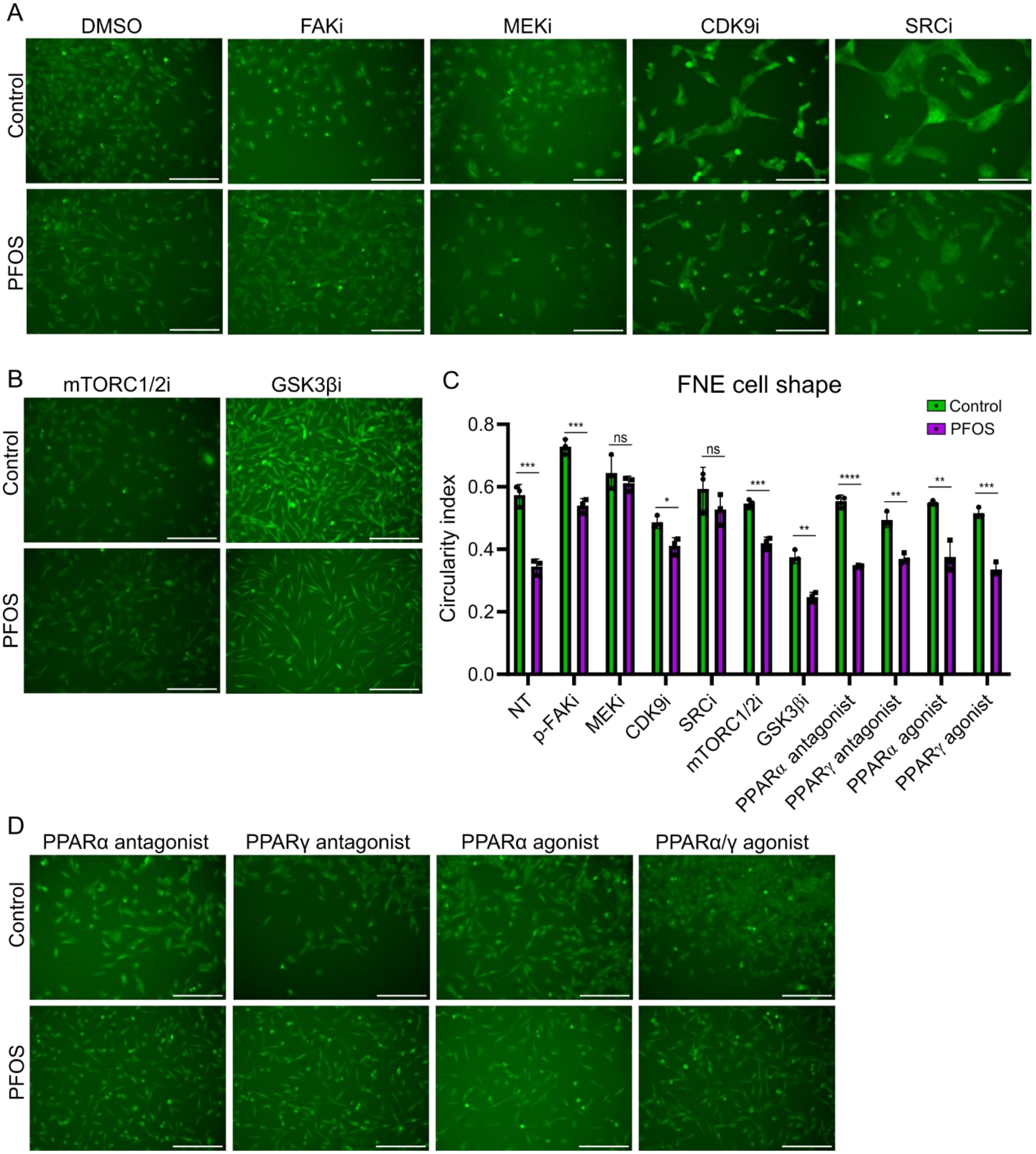


**Figure S5. Inhibition of MAPK signaling or cholesterol supplementation rescues PFOS-induced defects. (A)** Representative phase-contrast images of FNE cells treated with PFOS (25 μM, 48 h) ± pharmacologic inhibitors of MEK (PD0325901, 0.5 μM), FAK (PF-573228, 2 μM), Src (saracatinib, 5 μM), or CDK9 (dinaciclib, 50 nM). All inhibitors were added 24 h before PFOS. Scale bars = 50 μm. **(B)** Representative images of PFOS-treated FNE cells ± mTORC1/2 inhibitor

(Torin 1, 250 nM) or GSK3β inhibitor (CHIR99021, 3 μM). Scale bars = 50 μm. **(C)** Circularity index of GFP-expressing FNE cells treated with vehicle control (water) or PFOS (25 μM, 48 h) ± pharmacologic inhibitors of MEK (PD0325901, 0.5 μM), FAK (PF-573228, 2 μM), Src (saracatinib, 5 μM), or CDK9 (dinaciclib, 50 nM). All inhibitors were added 24 h before PFOS. NT = no inhibitor pre-treatment (DMSO as vehicle control). Each dot represents the mean of > 10 cells per replicate (n = 3). **(D)** Representative fluorescence images of FNE cells constitutively expressing GFP and exposed to 25 μM PFOS for 48 h ± PPAR pathway modulators. Cells were pre-treated for 24 h with 10 μM GW-6471 (PPARα antagonist), 10 μM GW-9662 (PPARγ antagonist), 10 μM WY-14643 (PPARα agonist), or 10 μM tesaglitazar (dual PPARα/γ agonist),. Scale bars = 50 μm. Statistical analysis by one-way ANOVA with Tukey's post-hoc test; *p < 0.05; **p < 0.01; ***p < 0.001; ****p < 0.0001.

**Movie 1**: **FNE control cell tracking.** Phase contrast time-lapse of live untreated (control) FNE cells followed over a period of 48 hours at 20-minute intervals to monitor their movement and proliferation. Time-lapse speed is 10 frames per second.

**Movie 2**: **PFOS-treated FNE cell tracking.** Phase contrast time-lapse of live PFOS-treated FNE cells followed over a period of 48 hours at 20-minute intervals to monitor their movement and proliferation. Time-lapse speed is 10 frames per second. PFOS was added 48 hours before the start of imaging.

**Movie 3**: **FNE control cell monolayer wound healing.** Phase contrast time-lapse of live untreated (control) FNE cell monolayer immediately after a scratch was introduced. FNE cell monolayer was followed over a period of 72 hours at 1-hour intervals to monitor gap closure. Movie speed is 15 frames per second.

**Movie 4**: **PFOS-treated FNE cell monolayer wound healing.** Phase contrast time-lapse of live PFOS-treated FNE cell monolayer immediately after a scratch was introduced. FNE cell monolayer was followed over a period of 72 hours at 1-hour intervals to monitor gap closure. Time-lapse speed is 15 frames per second. PFOS was added to the FNE cell monolayer 48 hours before the start of imaging.
